# NADH dehydrogenases contribute to extracellular electron transfer by *Shewanella oneidensis* MR-1 in bioelectrochemical systems

**DOI:** 10.1101/657668

**Authors:** Cody S. Madsen, Michaela A. TerAvest

**Affiliations:** Department of Biochemistry and Molecular Biology, Michigan State University, East Lansing, MI, USA; Department of Biomedical Engineering, Michigan State University, East Lansing, MI, USA

**Author notes:** corresponding author Address: 603 Wilson Rd., East Lansing, MI, 48823.

## Abstract

*Shewanella oneidensis* MR-1 is quickly becoming a synthetic biology workhorse for bioelectrochemical technologies due to a high level of understanding of its interaction with electrodes. Transmembrane electron transfer via the Mtr pathway has been well characterized, however, the role of NADH dehydrogenases in feeding electrons to Mtr has been only minimally studied in *S. oneidensis* MR-1. Four NADH dehydrogenases are encoded in the genome, suggesting significant metabolic flexibility in oxidizing NADH under a variety of conditions. Strains containing in-frame deletions of each of these dehydrogenases were grown in anodic bioelectrochemical systems with N-acetylglucosamine or D,L-lactate as the carbon source to determine impact on extracellular electron transfer. A strain lacking the two dehydrogenases essential for aerobic growth exhibited a severe growth defect with an anode (+0.4 V_SHE_) or Fe(III)-NTA as the terminal electron acceptor. Our study reveals that the same NADH dehydrogenase complexes are utilized under oxic conditions or with a high potential anode. Understanding the role of NADH in extracellular electron transfer may help improve biosensors and give insight into other applications for bioelectrochemical systems.

**TOC Graphic:** 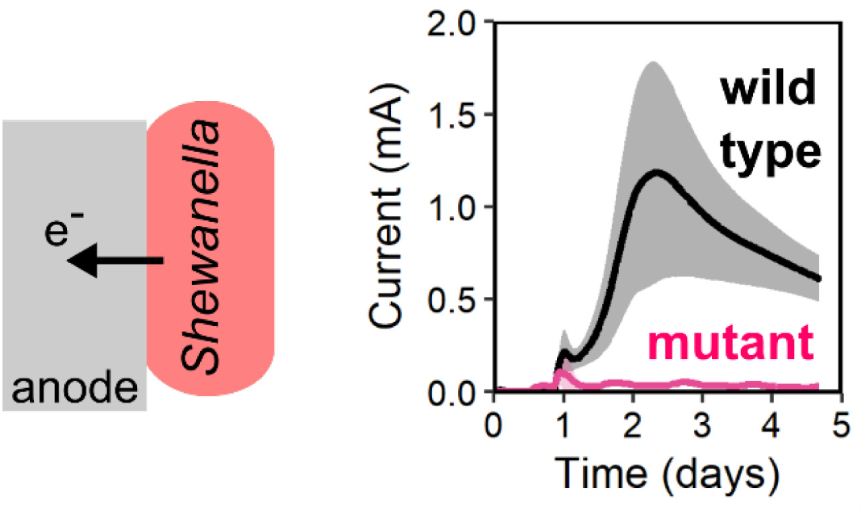

## Introduction

Bioelectrochemical systems (BESs) interface electrochemically active bacteria with electrodes for biotechnological applications including biosensing and electricity production.^1,2^ *Shewanella oneidensis* MR-1 is a γ-proteobacterium that has been used as a pure-culture model organism to study the biochemical underpinnings of microbe-electrode interaction in BESs. Detailed understanding of the Mtr electron transfer pathway in *S. oneidensis* MR-1 has enabled development of new bioelectrochemical technologies, such as genetically-encoded biosensors^3,4^ and strains with increased current production capability.^2,5^ Continued development of synthetic biology tools for electrochemically active bacteria, combined with increased knowledge of the molecular basis of extracellular electron transfer can increase functionality of bioelectrochemical technologies.^2,6^

The extracellular electron transfer capability of *S. oneidensis* MR-1 is dependent on the Mtr pathway,^7^ which is a porin-cytochrome complex (**Figure 1**).^8^ Porin-cytochrome complexes create transmembrane electron conduits that facilitate extracellular electron transfer.^9^ Once electrons reach the outer surface of the cell, multiple mechanisms are used to transfer electrons to solid terminal electron acceptors. The major mode of electron transfer in *S. oneidensis* relies on flavins as diffusible shuttles or bound cofactors to enhance electron transfer between an outer membrane cytochrome, MtrC, and solid minerals or electrodes.^7,10,11^ Flavins do not need to be added exogenously to enhance electron transfer rates because *S. oneidensis* MR-1 naturally secretes flavins in up to µM concentrations.^7,12^ MtrC can also transfer electrons directly to solid electron acceptors, although at a lower rate than flavin-based mechanisms.^10,12^

**Figure 1.**
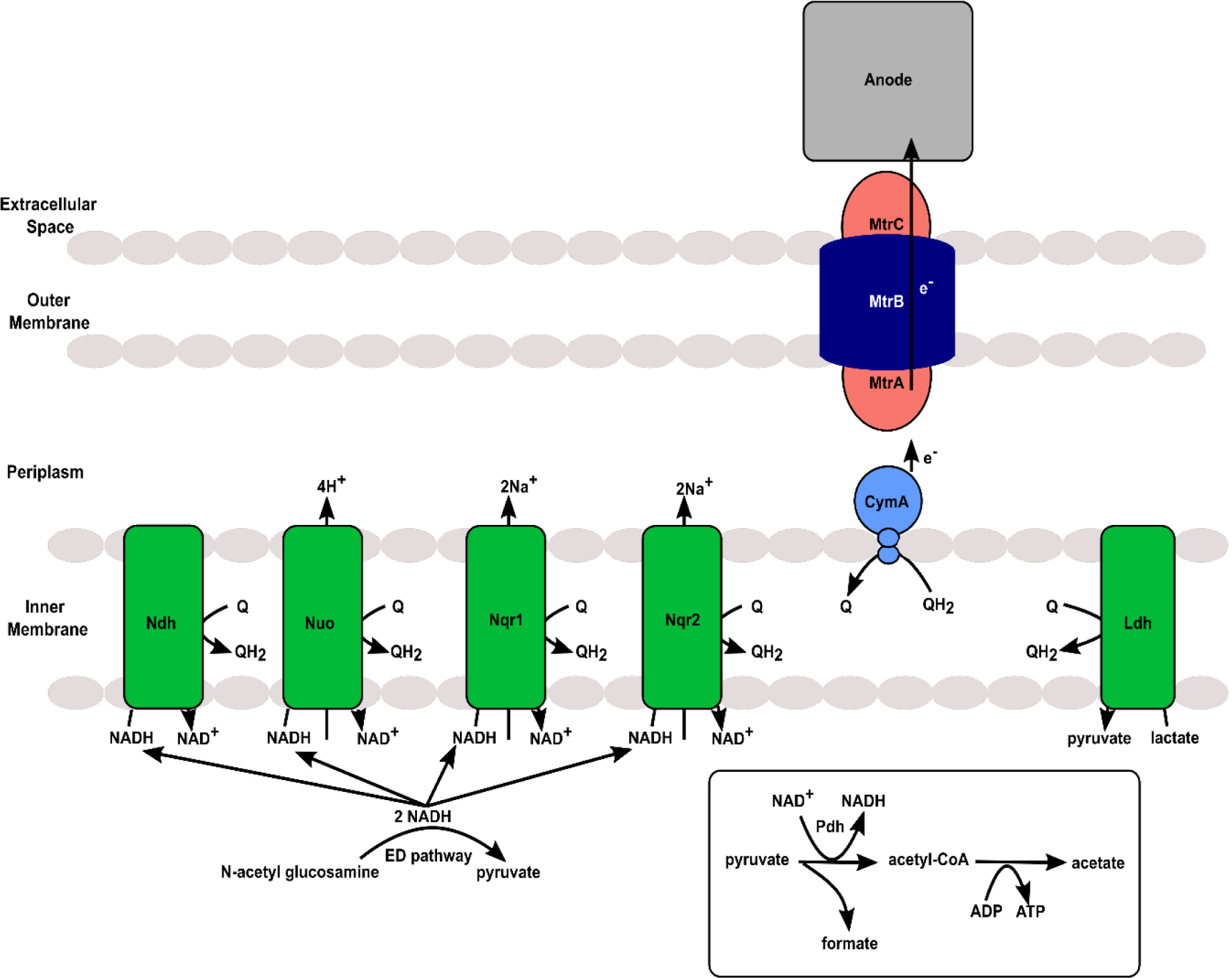
Metabolism of NAG and pyruvate and resulting electron transfer mechanisms within *Shewanella oneidensis* MR-1.

Electrons transported through the Mtr pathway are derived from upstream organic carbon metabolism and enter the inner membrane quinone pool via quinone-linked dehydrogenases, such as lactate dehydrogenase, or NAD^+^ -linked dehydrogenases coupled with NADH dehydrogenases (**Figure 1**). Hunt et al.^13^ proposed that anaerobic lactate oxidation by *S. oneidensis* MR-1 utilizes only quinone-linked dehydrogenases to bring electrons into the quinone pool, making NADH dehydrogenases theoretically unnecessary under these conditions. This model rests on the assumptions that pyruvate is processed only by pyruvate-formate lyase (Pfl) under anoxic conditions and that all acetyl-coA is converted to acetate and excreted. However, Hirose et al.^14^ recently demonstrated that the NAD^+^ -linked pyruvate dehydrogenase (Pdh) is activated under anoxic conditions with a high potential electron acceptor, and NADH dehydrogenase activity is necessary under such conditions. Beyond the importance of NADH dehydrogenases in anaerobic lactate metabolism, *S. oneidensis* also uses other substrates that must be oxidized by NAD^+^ -linked dehydrogenases, such as N-acetylglucosamine (NAG). NAG is a carbohydrate that is metabolized via the Entner-Doudoroff glycolysis pathway^15^ in *S. oneidensis* MR-1, resulting in production of NADH as an electron carrier.

Although Hirose et al.,^14^ demonstrated that NADH dehydrogenase activity is necessary for growth on lactate with high potential electrodes, their study did not investigate which specific NADH dehydrogenases are used. The *S. oneidensis* MR-1 genome encodes four putative NADH dehydrogenases: Nuo (SO_1009 to SO_1021), Ndh (SO_3517), Nqr1 (SO_1103 to SO_1108) and Nqr2 (SO_0902 to SO_0907) (**Figure 1**).^16,17^ Nuo couples NADH oxidation with proton translocation (4 H^+^ /2 e^−^), Nqr1 and Nqr2 act as sodium-ion translocators (2 Na^+^ /2 e^−^), and Ndh acts as a type II ‘uncoupling’ dehydrogenase that does not translocate ions.^16,17^ Ion translocation by these complexes generates a membrane potential that powers flagellar motors and ATP synthesis. Therefore, energy conservation from extracellular electron transfer will differ depending on which NADH dehydrogenases are active. We recently determined that activity of either Nuo or Nqr1 is essential for aerobic growth in minimal medium.^18^

To follow up on our previous study and work by Hirose et al.,^14^ we investigated the effects of deleting each NADH dehydrogenase on anaerobic respiration in *S. oneidensis*. We used mutant strains lacking single or multiple NADH dehydrogenases and studied their electron transfer capability in BESs under anoxic conditions.^18^ We utilized both NAG and D,L-lactate as substrates to investigate the effect of differing levels of NADH generation on phenotypic differences between the strains. Current production, growth, and major metabolites were compared between wild-type and all mutant strains. We learned that under the conditions tested here, *S. oneidensis* requires either Nuo or Nqr1, the same dehydrogenases needed under aerobic conditions.

## Materials and Methods

### Bacterial strains and culture conditions

*S. oneidensis* MR-1 cultures were inoculated from colonies on lysogeny broth (LB) agar plates and grown overnight in LB at 30°C with shaking at 275 rpm. After overnight growth, 1 mL was sub-cultured into 4 mL of fresh LB and grown for 8 hours (30°C, 275 rpm). NADH dehydrogenase mutants of *S. oneidensis* MR-1 (Δ*nuoN, Δndh, ΔnqrF1, ΔnqrF2, ΔnuoNΔnqrF1, ΔnqrF2Δndh, ΔnuoNΔndh*) were previously generated using an in-frame deletion method, and functions were confirmed via complementation studies.^18^ Knockout mutants were confirmed by PCR using flanking primers of the target knockout region.^18^

### Growth in bioelectrochemical systems

Two-chambered BESs were constructed as follows. The working and counter chambers were separated by a cation exchange membrane (Membranes International). M5 minimal medium^18^ was added to each working electrode chamber and 175 mL of PBS was added to each counter chamber. After autoclaving, Wolfe’s minerals and Wolfe’s vitamins^19^ were added to 1X concentration, riboflavin was added to 1 µM, and 20 mM D,L-lactate or 10 mM NAG were added, resulting in a final working anodic chamber volume of 160 mL. Working electrodes were constructed using carbon felt cut into 25 mm x 25 mm squares (Alfa Aesar, Part No: 43200). A segment of titanium wire (Malin Co., 0.025’’ titanium) was inserted through the carbon felt and sealed with carbon cement (Fluka Analytical, Part No: 09929-30G). Carbon rods, 1/8” in diameter (Electron Microscopy Science, 07200) were used for counter electrodes. Ag/AgCl/sat’d KCl reference electrodes were prepared in house. The assembled systems were placed on a magnetic stirrer and the electrodes were connected to a potentiostat (Bio-Logic Science Instruments). The working electrode potential was set to +0.2 V vs reference and current was measured every 1 second.

Baseline current measurements were collected for 24 hours before inoculation while bubbling nitrogen gas (Airgas, purity: 99.9%) into the working chamber. Each culture was pelleted, washed once with M5 minimal medium and standardized to an OD_600_ equal to 1.0. The BES working chambers were inoculated to an OD_600_ equal to 0.01 in a final volume of 160 mL. Samples for measurement of OD_600_ and substrate and product concentrations were taken daily. Samples were stored at -20°C prior to HPLC analysis. The working electrodes were removed at the end of the five day experiment for NAG and four day experiment for D,L-lactate and stored in 50 mL conical tubes at -20°C.

### Culturing *S. oneidensis* MR-1 with Fe(III)-NTA

M5/Fe(III)-NTA medium was prepared in 1 L volumes by preparing 450 mL of sterile 2X M5 minimal medium then adding 500 mL of filter sterilized (0.2 µm) 200 mM Fe(III)-NTA solution. The 200 mM Fe(III)-NTA medium was prepared by adding the following components to 400 mL of dH_2_O: 390 mM NaHCO_3_, 210 mM Fe(III)-NTA, 333 mM FeCl_3 20_ (Sigma Aldrich). The pH was adjusted to 6.8 using 5 M NaOH and the final volume was brought to 500 mL using dH_2_O. After combining the solutions, Wolfe’s minerals, Wolfe’s vitamins and 1 µM riboflavin were added. Medium and carbon sources were degassed in anaerobic chamber for five days prior to culture preparation. Cultures were inoculated to OD_600_=0.01 in a final volume of 5 mL and grown in an anaerobic chamber without shaking at 30°C with 10 mM D,L-lactate or NAG. The cultures were sampled for reduction of Fe^3+^ daily by diluting 25 µL of culture with 75 µL of degassed 2 M HCl in an anaerobic chamber and storing the samples at -20°C. Culture samples of 1 mL were taken at the beginning and end of the experiment and stored at -20°C. NAG, lactate, and acetate concentrations were determined by HPLC as described previously.^18^

### Measuring Fe^3+^ reduction

Fe^2+^ standards were prepared anaerobically at concentrations of 5, 10, 25, 50, 75, 100 and 125 mM. Standards and samples were diluted 1:20 in 0.5 M HCl before adding ferrozine reagent. Ferrozine reagent was prepared by adding 1 g/L ferrozine to 50 mM (11.9 g/L) HEPES and pH was adjusted to 7.0.^21^ The diluted standards and samples from each culture were added to a 96 well plate (VWR, Part No: 734-2327) in 10 µL aliquots and mixed with 190 µL of ferrozine reagent.^21^ Absorbance was read at 562 nm (A_562_) in a 96-well plate reader (SpectraMax, Part No: M2).^21^ Standard curves were prepared by graphing concentration of Fe^2+^ (mM) versus A_562_ to convert absorbance of samples to concentrations.

### Total protein analysis

Samples from BESs were spun down at 13,000 x g for 10 minutes and the supernatant was removed. The pellets were resuspended in 1 mL of 0.2 M NaOH and heated at 95°C for 30 minutes with mixing every 10 minutes. The procedure was repeated with working electrodes by suspending them in 15 mL of 0.2 M NaOH then vortexing, heating and mixing them. A Pierce BCA Protein Assay Kit (Thermo Scientific) was used for measurement of total protein.

### Data analysis

Current, OD_600_, HPLC, Fe^2+^ and BCA protein measurements were evaluated and plotted using Rstudio using the following packages: ggplot2,^22^ reshape2,^23^ dplyr,^24^ plyr,^25^ and TTR.^26^

## Results and discussion

### Nuo or Nqr1 is required for growth with an electrode or Fe(III)-NTA

In-frame deletions of each of the four NADH dehydrogenases in *S. oneidensis* MR-1 were generated in a previous study (**Figure 1**).^18^ The present study investigates single-knockouts of each NADH dehydrogenase (Δ*nuoN*, Δ*nqrF1*, Δ*nqrF2*, and Δ*ndh*), and a subset of the possible double knockout combinations (Δ*nuoN*Δ*nqrF1*, Δ*nuoN*Δ*ndh*, and Δ*nqrF2*Δ*ndh*). The wild-type (WT) and mutant strains were grown in BESs with 10 mM NAG or 20 mM D,L-lactate and an electrode potential of +0.4 V_SHE_. Under these conditions, none of the single mutant strains showed a significant difference from WT in electric current production, growth, or substrate utilization (**Figures 2 and S1**). Of the double knockout strains, only Δ*nuoN*Δ*nqrF1* performed significantly differently from WT. The Δ*nuoN*Δ*nqrF1* strain produced 35-fold less current and 11-fold less planktonic biomass than WT. The Δ*nuoN*Δ*nqrF1* strain consumed essentially no NAG (**Figure 2A-C**). When grown with D,L-lactate, the Δ*nuoN*Δ*nqrF1* strain produced 19-fold less current and 12-fold less planktonic biomass compared to WT without metabolizing a significant amount of D,L-lactate by the end of the experiment (**Figure 2D-F and S2**).

**Figure 2.**
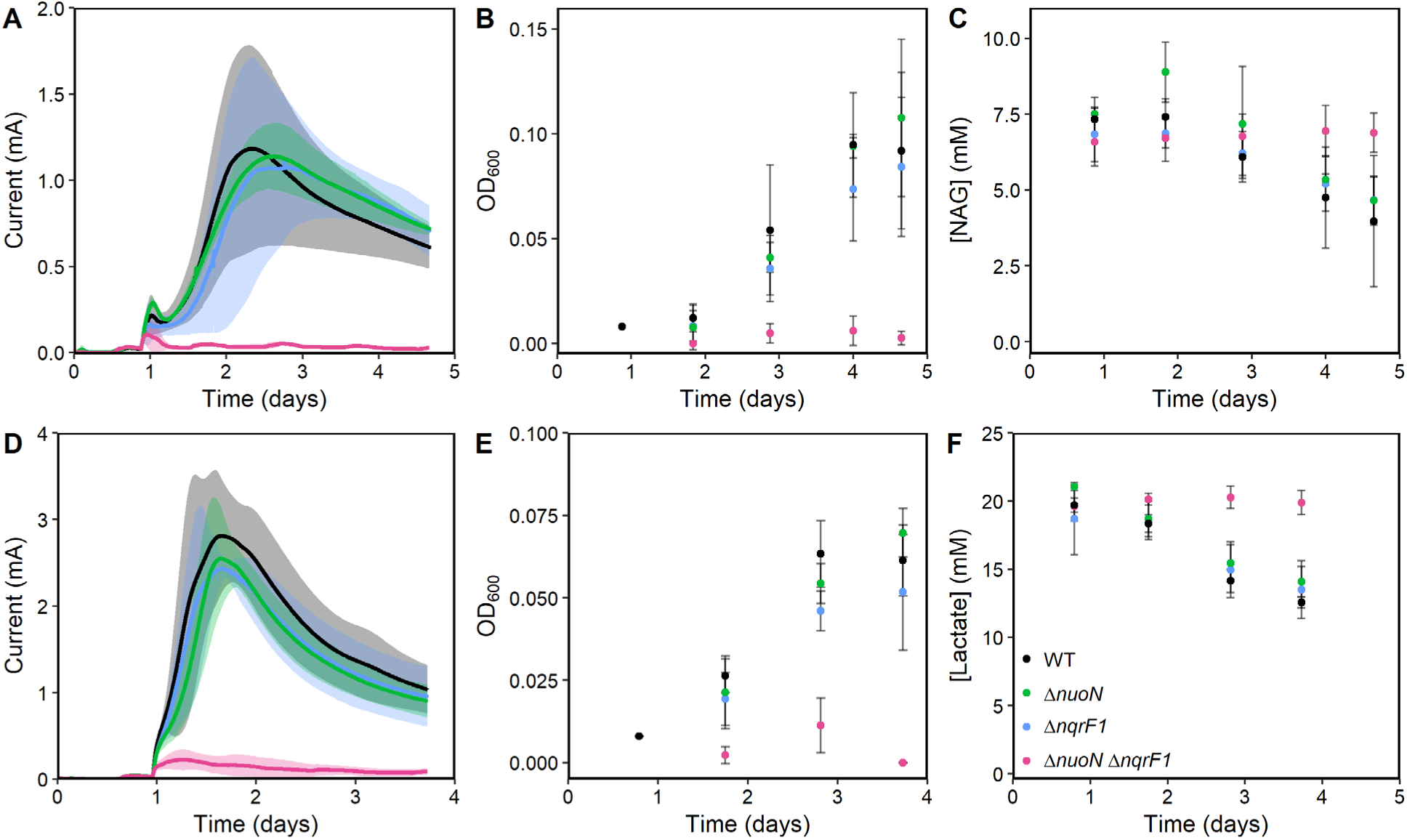
Current production (A), OD_600_ (B) and NAG metabolism (C) by WT, Δ*nuoN*Δ*nqrF1* and single mutants with 10 mM NAG as carbon source. Current production (D), OD_600_ (E) and D,L-lactate metabolism (F) by WT, Δ*nuoN*Δ*nqrF1* and single mutants with 20 mM D,L-lactate as carbon source. Points or lines represent an average of multiple replicates and error bars represent standard deviation (n=3).

We also conducted experiments using Fe(III)-NTA as the electron acceptor in an anaerobic chamber. Fe(III)-NTA has a redox potential of +0.587 V_SHE_ at pH 7, similar to the electrode potential of +0.4 V_SHE_. Under these conditions, all single knockout strains again grew similarly to WT (**Figure 3**). Only the Δ*nuoN*Δ*nqrF1* showed a significant growth defect, producing 6.4 ± 0.73 mM and 6.1 ± 0.13 mM of Fe^2+^ when using NAG or D,L-lactate respectively, compared 26 ± 2.2 mM Fe^2+^ and 88 ± 7.0 mM Fe^2+^ for WT, respectively (**Figure 3**). The Δ*nuoN*Δ*nqrF1* strain also consumed minimal NAG or D,L-lactate during iron reduction (**Figure 3**).

**Figure 3.**
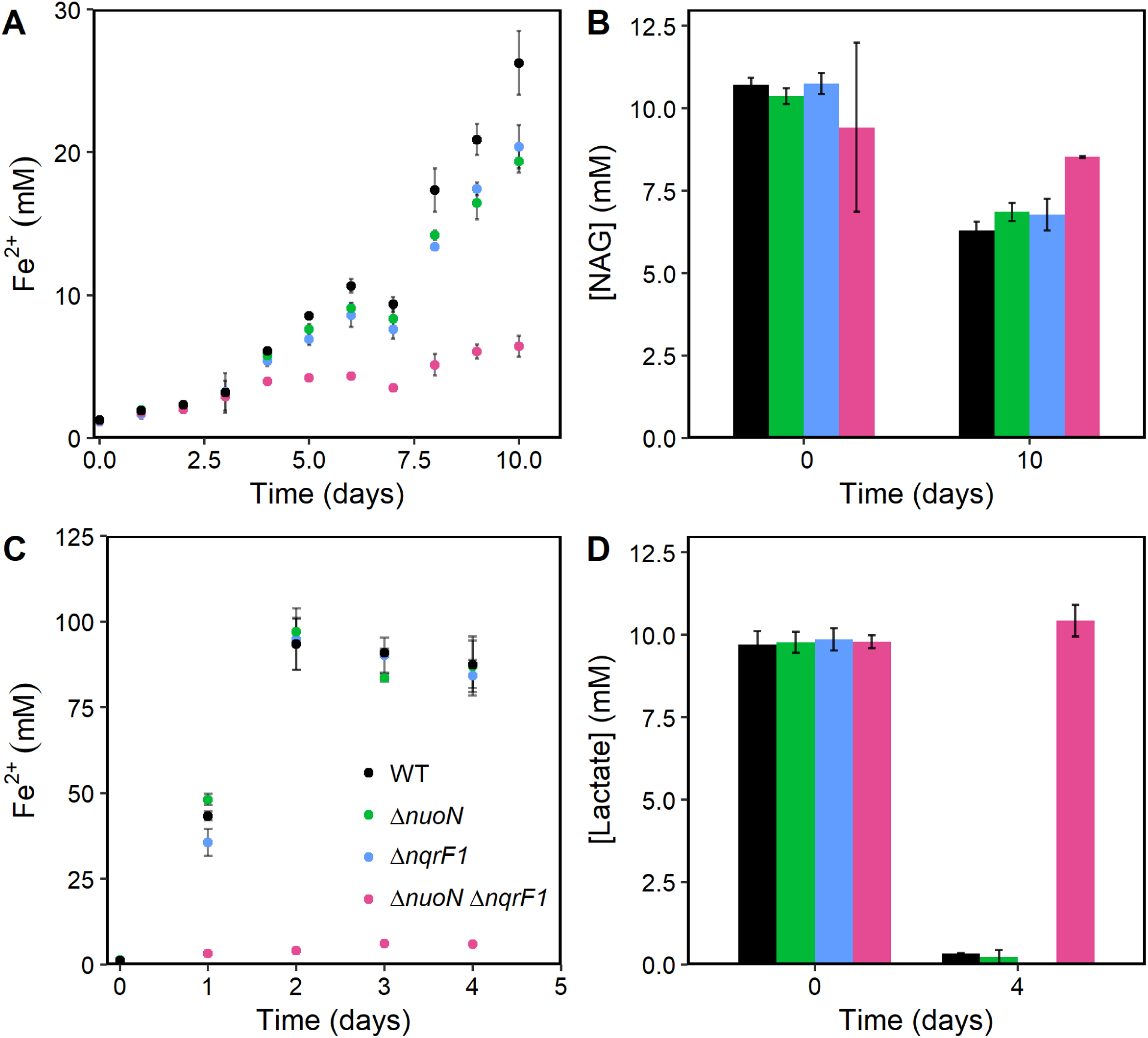
Reduction of Fe^3+^ (A) and NAG metabolism (B) by WT, Δ*nuoN*Δ*nqrF1* and single mutants with 10 mM NAG as the carbon source. Reduction of Fe^3+^ by (C) and D,L-lactate metabolism (D) by WT, Δ*nuoN*Δ*nqrF1* and single mutants with 10 mM D,L-lactate as the carbon source. Points represent an average of multiple replicates and error bars represent standard deviation (n=3).

### NADH dehydrogenase usage is similar between aerobic and anaerobic growth

We initially hypothesized that NADH dehydrogenase mutants would show no growth defect in BESs with D,L-lactate based on previous predictions that NADH is not an electron carrier in anaerobic lactate metabolism by *S. oneidensis* MR-1.^16,17^ However, we observed that either Nuo or Nqr1 was necessary for growth under these conditions. This can be explained by anaerobic Pdh activity, which generates NADH during pyruvate oxidation. Hirose et al.^14^ recently observed that Pdh is transcriptionally upregulated at high electrode potentials or with high potential minerals. We used electron acceptors with high potentials in this study (electrode at +0.4 V_SHE_ and Fe(III)-NTA at E°= +0.587_SHE_), therefore Pdh was likely upregulated and active in our system. It appears that Pdh was the main source of NADH, rather than the TCA cycle, because most acetyl-coA was converted to acetate and excreted. We observed production of ∼2.5 moles of acetate for every mole of NAG consumed (3 is the theoretical maximum), and ∼1.3 moles of acetate per mole of lactate consumed (1 is the theoretical maximum).

Surprisingly, we observed that the same two NADH dehydrogenases necessary for aerobic growth, Nuo and Nqr1, were necessary for anaerobic growth with high potential electron acceptors. Previous high-throughput fitness analysis suggested that Nqr2 and Ndh would be more important for growth under anoxic conditions.^18,27^ However, these two complexes were not sufficient to maintain growth under the conditions tested here. Considering that oxygen is also a high-potential electron acceptor, our results suggest that regulation of the NADH dehydrogenases may be more sensitive to the redox potential of the electron acceptor rather than its specific identity. Indeed, Hirose et al.^14^ found that Nuo transcript levels were higher with a high-potential electrode than with a low-potential electrode. Our results confirm that Nuo is important for respiration under very similar conditions. Because Nuo is the most efficient of the four NADH dehydrogenases, its upregulation with high potential electron acceptors may allow *S. oneidensis* MR-1 to generate more ATP by harnessing the greater voltage gap between quinols and the terminal electron acceptor. The capability of *S. oneidensis* MR-1 to reshape its electron transport chain to maximize energy production with a variety of electron acceptors likely has significant impacts on its growth in different conditions.

## Conclusions

We assessed the impact of NADH dehydrogenase deletions on current production, growth and metabolism in *S. oneidensis* MR-1. Our results suggest that Nuo and Nqr1 are both involved in electron transfer from NADH to the Mtr pathway and at least one is necessary for growth with high-potential anodes or Fe(III)-NTA as the electron acceptor. These results indicate that neither Nqr2 nor Ndh are sufficient to maintain electron flux to the Mtr pathway under these conditions. This fundamental science may unlock potential applications in BESs and biosensors and facilitate understanding of electron transfer in *S. oneidensis* MR-1.

## Acknowledgements

The authors thank Nicholas Tefft for assisting in bioreactor sampling and HPLC analysis, Kody Duhl for developing *S. oneidensis* mutants, and Dr. Magdalena Felczak for insightful questions on the project. The authors also thank Dr. Neal Hammer for helpful comments on the manuscript. Support for this work was provided by internal funding to the PI from Michigan State University, and the Richard L. Anderson Award and the Pamela Fraker Research Scholarship to Cody Madsen. The Michigan State University College of Natural Science and Department of Biochemistry and Molecular Biology also provided six research stipends to support this work. This work was also supported by the USDA National Institute of Food and Agriculture, Hatch project 1009805.

