## Supplemental Figures for "NADH dehydrogenases contribute to extracellular electron transfer by *Shewanella oneidensis* MR-1 in bioelectrochemical systems"

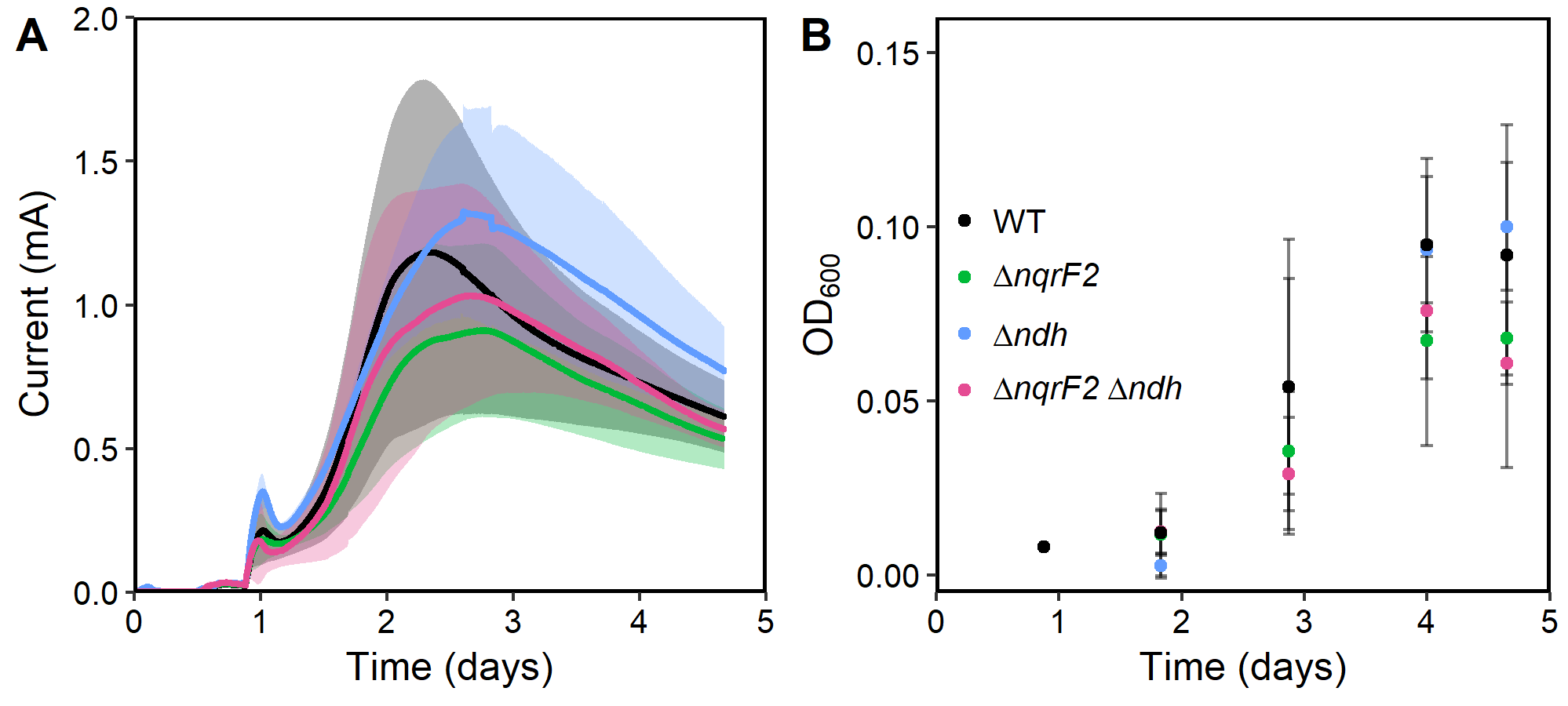

**Figure S1.** Current production (A) and OD_600_ (B) by WT, ∆*nqrF2*∆*ndh* and correlating single mutants with 10 mM NAG as the carbon source. Data analyzed in R with shaded regions and error bars indicating standard deviations (n=3).

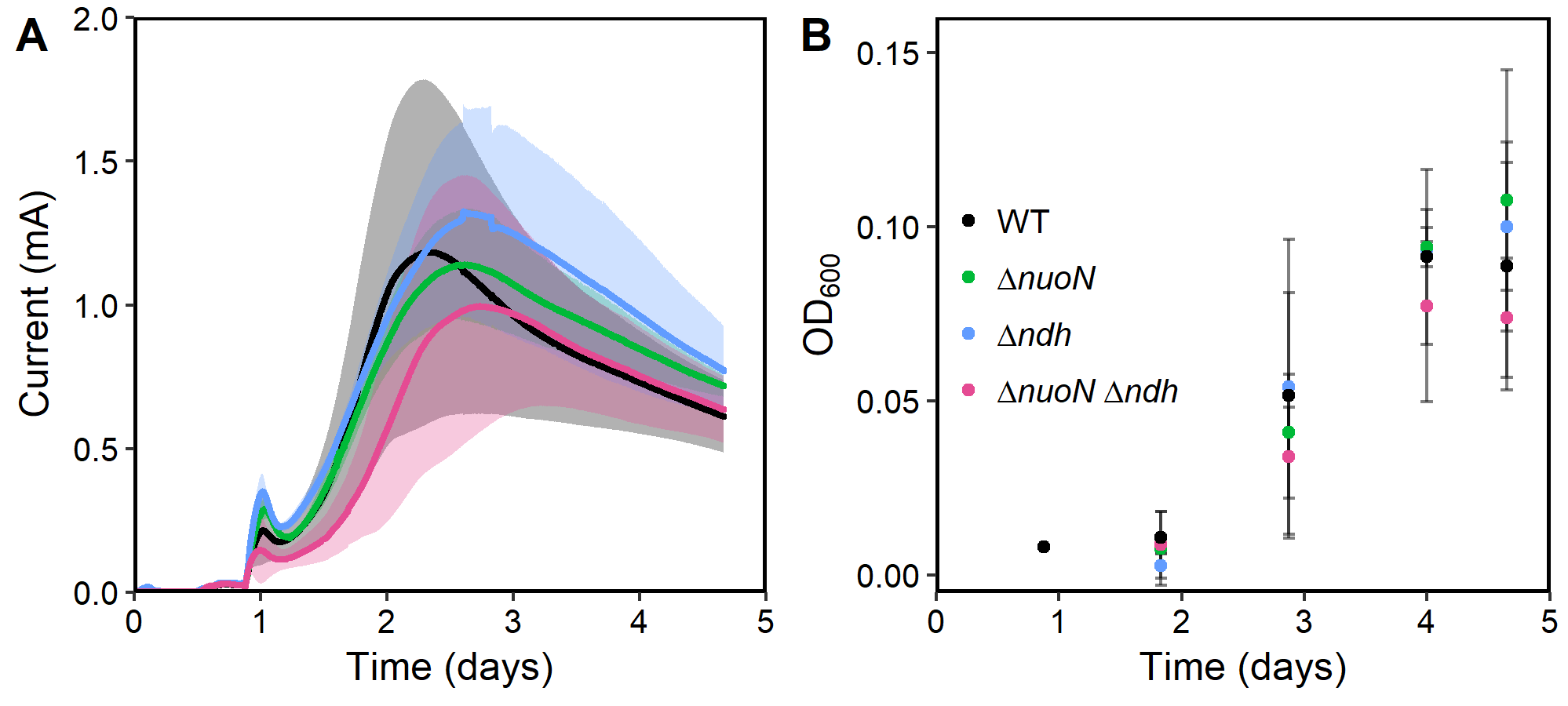

**Figure S2.** Current production (A) and OD_600_ (B) by WT, ∆*nuoN*∆*ndh* and correlating single mutants with 10 mM NAG as the carbon source. Data analyzed in R with shaded regions and error bars indicating standard deviations (n=3).
